# Phase-Restoring Subpixel Image Registration: Enhancing Motion Detection Performance in Fourier-domain Optical Coherence Tomography

**DOI:** 10.1101/2022.06.15.496241

**Authors:** Huakun Li, Bingyao Tan, Vimal Prabhu Pandiyan, Veluchamy Amutha Barathi, Ramkumar Sabesan, Leopold Schmetterer, Tong Ling

## Abstract

Phase-sensitive Fourier-domain optical coherence tomography (FD-OCT) enables in-vivo, label-free imaging of cellular movements with detection sensitivity down to the nanometer scale, and it is widely employed in emerging functional imaging modalities, such as optoretinography (ORG), Doppler OCT, and optical coherence elastography. However, when imaging tissue dynamics in vivo, tissue movement or bulk motion introduces decorrelation noise that compromises motion detection performance, particularly in terms of sensitivity and accuracy. Here, we demonstrate that the motion-related decorrelation noise in FD-OCT can be accurately corrected by restoring the initial sampling points using our proposed Phase-Restoring Subpixel Image Registration (PRESIR) method. Derived from a general FD-OCT model, the PRESIR method enables translational shifting of complex-valued OCT images over arbitrary displacements with subpixel precision, while accurately restoring phase components. Unlike conventional approaches that shift OCT images either in the spatial domain at the pixel level or in the spatial frequency domain for subpixel correction, our method reconstructs OCT images by correcting axial displacement in the spectral domain (k domain) and lateral displacement in the spatial frequency domain. We validated the PRESIR method through simulations, phantom experiments, and in-vivo optoretinography in both rodents and human subjects. Our approach significantly reduced decorrelation noise during the imaging of moving samples, achieving phase sensitivity close to the fundamental limit determined by the signal-to-noise ratio (SNR).

## 1. Introduction

Fourier-domain optical coherence tomography (FD-OCT) enables depth-resolved imaging via the Fourier transform of the spectral interferogram between the back-scattered light from the sample and the reference beam [1]. While conventional OCTs utilize the amplitude component of OCT signals, which represents the light intensity scattered from anatomical microstructures, phase-sensitive OCT exploits the phase difference between repeated A-scans [2], B-scans [3, 4], or volumetric scans [5] to achieve imaging of structural dynamics with sensitivity down to the nanometer scale [2]. Owing to its high motion sensitivity, phase-sensitive OCT has led to the development of several label-free functional imaging modalities. For instance, Doppler OCT has been widely used in velocimetry to quantify blood flow velocity [6], while optical coherence elastography was developed to evaluate tissue biomechanical properties [4, 7]. Recently, the application of phase-sensitive OCT for imaging the functional activity of photoreceptors in response to light stimuli, generally termed optoretinography (ORG), has sparked significant interest in both scientific research and clinical diagnosis [8, 9, 10, 11]. Furthermore, phase-sensitive OCT may pave the way for all-optical interferometric thermometry in non-damaging photothermal therapies [12, 13].

Despite the ever-growing impact of phase-sensitive OCT, it is well known that the measured phase change possesses a decorrelation noise besides the desired optical path difference (OPD) and the signal-to-noise ratio (SNR) dependent phase noise. Even without any change in the tissue structure that causes the speckle pattern to decorrelate, the decorrelation noise still occurs when the sample undergoes bulk movements. Such motion-related decorrelation noise is introduced by the mismatch between the sampling points in consecutive OCT images [14, 15], which can be problematic for in-vivo measurements where the sample is constantly affected by vascular pulsation, breathing, and other involuntary movements. When the bulk movement of the sample is of interest, the decorrelation noise can severely degrade the fidelity of the measured axial movement, especially if the magnitude of the movement goes near or above the OCT system’s axial resolution (on the order of a few microns) [16]. Alternatively, when examining local tissue deformation (e.g. for in-vivo optoretinography), it is common practice to use the self-referenced method to cancel out unwanted phase drifts caused by the bulk tissue movement and the inevitable fluctuations in the OPD between the sample arm and the reference arm [17]. However, the detrimental effect of decorrelation noise in this scenario has long been overlooked, as the phase uncertainty due to the sampling point mismatch can still vary from one pixel to another along the same A-line [6].

In fact, the complex-valued OCT signal at each pixel can be modeled as a coherent superposition of scattered light from adjacent scatterers, weighted by the point spread function (PSF) centered at that pixel [18, 19]. As illustrated by the numerical simulations in a recent report by Hepburn et al. [20], the phase inaccuracy in speckles with respect to the actual motion was seen to result from the change in the weights of scatterers in the aforementioned superposition. A corollary of this finding is that any tissue movement will change the positions of individual scatterers relative to the pixel of interest in the OCT image, inevitably altering their scattered light’s contribution to the coherent superposition and leading to an unwanted phase variation. From this perspective, the extra phase variation due to the sampling point mismatch can be regarded as a deterministic error (termed “motion-induced phase error” in this article) instead of stochastic noise, if the scatterers’ relative locations remain fixed in the sample [21]. In other words, maintaining identical sampling points during the sample movement, which can be achieved by post-hoc image registration, may allow mitigating or even eliminating the motion-related decorrelation noise entirely. Moreover, the effectiveness of such an approach depends on the accuracy with which we can restore the original sampling points, preferably down to the subpixel level.

Typically, image registration in OCT involves two steps: motion estimation and image correction. Although normalized cross-correlation (NCC)-based [22, 23] and phase-only correlation-based methods [24] enable subpixel level motion estimation in both axial and lateral directions, conventional image correction approaches, which shift the original complex-valued OCT image [25, 26] or the 2-4 fold upsampled OCT image [27] in pixels, are limited by the discrete interval determined by the upsampling rate. To facilitate image shifting over arbitrary distances, an intuitive thought (referred to as the FT-based method in this article) is to treat complex-valued OCT images as digital images and directly multiply the Fourier transform (FT) of the image by an exponential term [28]. However, Lee et al. reported that the FT-based method’s performance, evaluated by the magnitude of cross-correlation between registered images, was significantly inferior to that of upsampling methods [28], likely due to the omission of the unique physics underlying the axial formation of the FD-OCT signal.

In this article, we propose a Phase-Restoring Subpixel Image Registration (PRESIR) method that takes into account the nuances between the FD-OCT signals in the axial and lateral directions. Our proposed method is capable of accurately restoring phase components when shifting images in FD-OCT. In addition, we identified the root cause of the motion-induced phase error using analytic formulas and analyzed its influence on motion detection accuracy through numerical simulations. Notably, we found that the PRESIR method eliminated the motion-induced phase error and achieved motion detection sensitivity approaching the theoretical limit in simulations and synthetic phantom experiments. Moreover, we compared the motion detection sensitivity of the pixel-level image registration method, the FT-based method, and the proposed PRESIR method when detecting nanoscopic tissue deformations within moving samples. We found that the PRESIR method significantly improved detection performance in in-vivo ORG for rodents and human subjects.

This work significantly expands upon our preliminary findings presented at the SPIE Photonics West 2023 [29, 30] by incorporating more rigorous analyses of the proposed methodology and thorough experimental validations. Besides, no conference paper was published alongside those previous oral presentations.

## 2. Methods

### 2.1 Analytic explanation of the motion-induced phase error in FD-OCT

OCT signals originate from the light scattered from individual scatterers within a sample [18]. If the sample is physically displaced by Δ*x* and Δ*z* in the lateral and axial directions, respectively, the reflectivity distributions *η*_*R*_(*x*, *z*) and *η*_*T*_(*x*, *z*) of the reference frame and target frame can be written as,

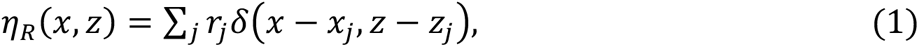

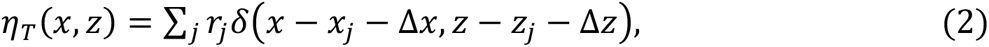

where *δ*() is the Dirac delta function, *j* is the index of scatterers, *r*_*j*_ and (*x*_*j*_, *z*_*j*_) denote the electric field reflectivity and the coordinate of the *j*^th^ scatterer in the reference frame.

Taking into account Eqs. (1-2) and the general FD-OCT model (see Appendix A), we can derive the complex-valued OCT images 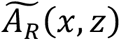 and 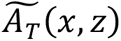 measured from the reference and target positions as follows:

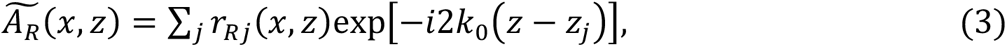

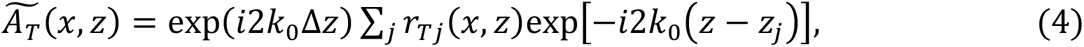

where *r*_*R*_*_*j*_*(*x*, *z*) and *r*_*T*_*_*j*_*(*x*, *z*) represent the PSF-weighted reflectivities of the *j*^th^ scatterer in the reference and target frames, respectively, as defined in Appendix B.

Consequently, we can express the phase difference Δ*φ* between the complex-valued OCT image 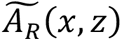 and 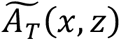 as follows:

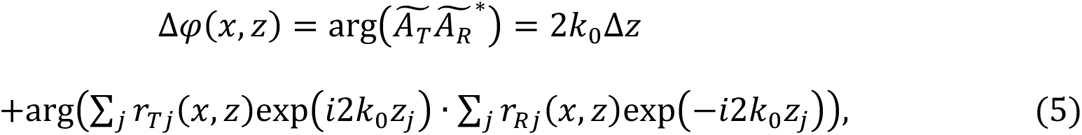

where arg() denotes the calculation of argument, and * is the complex conjugate operation.

Equation (5) demonstrates that the phase difference Δ*φ*(*x*, *z*) is affected not only by a phase change of 2*k*_0_Δ*z*, which directly corresponds to the desired OPL change, but also by an additional error term arising from the variation in the PSF-weighted reflectivity that occurs with sample movement. Throughout this article, we denote this error as the motion-induced phase error. While our derivation is limited to motion within the (*x*, *z*) plane, it is important to note that the motion-induced phase error is also applicable to out-of-plane movement along the *y*-axis. Intriguingly, the above derivation reveals that the motion-induced phase error is pixel-specific and varies from pixel to pixel. Hence, self-referenced measurements that determine the phase difference between two pixels at different depths are ineffective in eliminating the motion-induced phase error.

### 2.2 Phase-restoring subpixel image shifting for accurate motion correction in FD-OCT

We conceived our phase-restoring subpixel image shifting approach from the general FD-OCT model (see Appendix A), where the way FD-OCT signals are formed in the axial direction departs fundamentally from how digital images are captured by standard cameras. As translational motion between cross-sectional frames is the dominant source of motion artifacts in high-speed OCT imaging, compared with rotational movement or image distortion within individual cross-sections [26], this study mainly concerns and corrects the translational shifts between the reference and target images.

We assume that the measured sample undergoes a translational shift of Δ*x* in the lateral (*x*) direction and Δ*z* in the axial (*z*) direction from the reference frame to the target frame. According to the general FD-OCT model (see Eq. (A4) in Appendix A), the complex-valued spectral interferograms of the reference frame 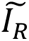 and the target frame 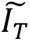 can be modeled as,

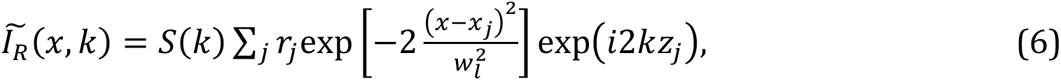

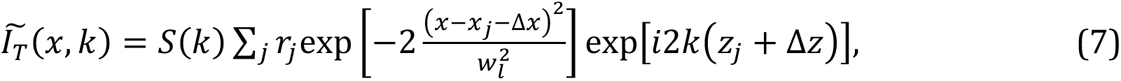

where *k* is the wavenumber, *S*(*k*) is the power spectrum of the light source, *r_j_* and (*x*_*j*_, *z*_*j*_) denote the electric field reflectivity and the coordinate of the *j*^th^ scatterer in the reference frame, *w*_*l*_is the 1⁄e^2^ spot radius of the OCT beam focused on the sample. Note that the axial coordinate (*z*) in this study, as well as *z*_*j*_ and Δ*z* mentioned earlier, represents the optical path length (OPL) resulting from both refractive index and axial location of the sample.

From Eqs. (6) and (7), it is evident that the axial location of an individual scatterer results in an exponential term exp(*i*2*kz*_*j*_) in the detected OCT signal, where 2*kz*_*j*_is the OPD between the beam scattered from the sample and the reference beam. In contrast, the OCT signal in the lateral direction is determined by the scatterer’s lateral location convolved with a PSF, similar to capturing digital images using standard cameras. Considering the difference in the imaging principle in the axial direction, if we manually multiply the spectral interferogram in FD-OCT with a numerical term exp(*i*2*k*Δ*z*′), we can arbitrarily shift the image axially by any displacement with an OPL of Δ*z*′ as if the sample were displaced by the exact same distance in the real world. Unlike conventional subpixel image shifting approaches, the above process does not require interpolation or a sampling frequency that is sufficiently high to avoid aliasing.

More specifically, comparing Eqs. (6) and (7), the spectral interferograms before and after the displacement satisfy the following relation [31, 32],

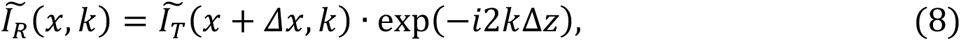

and it can be further written as,

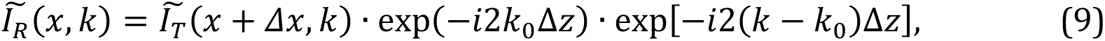

where *k*_0_is the center wavenumber of the light source spectrum. The first exponential term exp(−*i*2*k*_0_Δ*z*) accounts for the change of OPL, and the second wavenumber-dependent term exp[−*i*2(*k* – *k*_0_)Δ*z*], in fact, corresponds to the sampling point mismatch induced by the axial motion.

When applying the above findings to image registration in FD-OCT, we found that multiplying both aforementioned exponential terms enables accurate reconstruction of complex-valued OCT images as if the sample were physically shifted back, as written in Eq. (10),

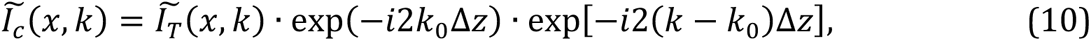

where 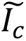 is the accurately reconstructed spectral interferogram after the axial correction. Equation (10) is useful for eliminating the negative influence of bulk motion in self-referenced measurements, such as when imaging local tissue deformations in vivo.

Notably, we can also correct the OCT image by multiplying only the second term, which eliminates the extra motion-induced phase error while retaining the OPL change, as shown in Eq. (11),

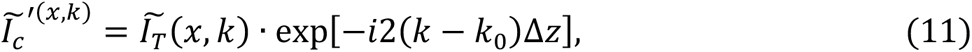

where 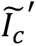 is the corrected spectral interferogram that retains the OPL change. When using phase signals to detect the sample’s bulk movement, Eq. (11) can help isolate and eliminate the motion-induced phase error that is coupled in the phase signal, allowing us to obtain the desired OPL change.

To effectively correct for lateral motion, the sample must be sufficiently oversampled in the lateral direction. Besides, the extra phase noise between adjacent A-scans, caused by bulk sample motion and system instabilities (e.g., galvanometer jitter and reference arm fluctuations) [33], must be significantly below *π* to maintain a stable phase relationship. Under these prerequisites, our lateral bulk motion correction strategy is similar to the FT-based method, where we conduct the image shift by multiplying an exponential term in the spatial frequency domain [28, 34]:

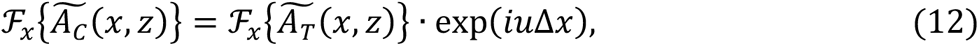

where 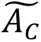 is the reconstructed complex-valued OCT image after lateral correction, and *u* represents the lateral spatial frequency. Note that (Δ*x*, Δ*z*) refers to the displacement from the reference frame to the target frame. When performing motion correction using Eqs. (10)-(12), the target frame should be translated back by (−Δ*x*, −Δ*z*) to its original position.

When benchmarking our proposed method, we compared its performance with the pixel-level correction and FT-based subpixel motion correction techniques. We also turned off the piezo actuator in phantom experiments to provide references for imaging static samples. In the FT-based method, complex-valued OCT images were treated as digital images, and axial and lateral displacements were corrected by multiplying the spatial frequency domain with exp(*iu*Δ*x*) · exp(*iv*Δ*z*), where *u* and *v* denote the lateral and axial spatial frequencies, respectively [28]. For pixel-level correction, we rounded the subpixel bulk movement, estimated using the methods in the next section, to the nearest integer. Motion correction was then performed by directly shifting the image by the negative value of the rounded integer.

### 2.3 Subpixel motion estimation from repeated cross-sectional scans and repeated volumetric scans

We used various existing methods to achieve motion estimation down to the subpixel level. In phantom experiments involving only axial motion, we estimated the subpixel-level axial bulk motion using the phase change extracted from the phantom surface.

For rodent retinal imaging and phantom experiments with two-dimensional movements in the axial and lateral directions, we adopted the single-step DFT approach [22, 23] to estimate subpixel-level displacements between the first and subsequent cross-sectional B-scans. Specifically, a 2-fold upsampled NCC function was first obtained by zero-padding the Fourier spectrum. The location of its peak was found as the initial estimation. Since only a small neighborhood around the NCC peak is of interest, the matrix-multiply DFT method was used to compute the *κ*-fold upsampled NCC map in a 1.5 x 1.5 pixel neighborhood centered on the initial estimation [23]. In this study, *κ* was set to 1000 to allow motion estimation with sufficiently small intervals. Compared with conventional upsampling strategy using fast Fourier transform (FFT), the matrix multiplication approach can greatly reduce the computational load.

In repeated volumetric scans for human retinal imaging experiments, we estimated the translational motion of individual B-scans with respect to the B-scans in a pre-selected reference volume. We followed the coarse-to-fine strategy proposed by Do et al. [25, 26], except for that in the fine estimation step, we adopted the single-step DFT approach to extend the previous pixel-level estimation to subpixel-level in the depth (*z*) and the line (*x*) dimensions. Briefly, during the coarse estimation step, we first selected a set of sub-volumes containing consecutive B-scans from the target volume. We then estimated the positions of these sub-volumes in the reference volume using the three-dimensional NCC method. Subsequently, for each B-scan in the target volume (referred to as the target B-scan), we determined its coarse shift in the scan (*y*) direction by linearly interpolating the sparse shifts of the sub-volumes. In the fine estimation step, for each target B-scan, we selected a sub-volume from the reference volume based on the interpolated coarse shift in the scan (*y*) direction. We then applied the single-step DFT approach to calculate the correlation value and the associated subpixel-level translational motion between the target B-scan and each B-scan in the selected reference sub-volume. Lastly, for each target B-scan, we obtained the estimated shifts (Δ*x*, Δ*y*, Δ*z*) in the line, scan, and depth directions that produced the highest correlation value.

### 2.4 Evaluation metrics for quantitative assessment of motion correction performance

We validated the motion-induced phase error and assessed the performance of motion correction techniques on moving phantom samples and in-vivo retinas. Samples without local deformation allow us to quantitatively characterize the accuracy and sensitivity of phase signals in OCT images: Accuracy, which is defined as the deviation of phase signals from the ground truth movement, was estimated by calculating the spatial standard deviation (*σ*_*s*_) of the phase signals across all pixels at a single time point; Sensitivity, on the other hand, corresponds to the phase fluctuation over time and was evaluated by computing the temporal standard deviation (*σ*_*t*_) of phase variation across all time points. To provide a more intuitive understanding of how these metrics affect motion detection accuracy and sensitivity, we converted the phase units (radians) to their corresponding OPL using the equation OPL = *φ*/2*k*_0_, where *φ* is the phase and *k*_0_ is the center wavenumber of the light source spectrum.

## 3. Experiments

### 3.1 Point-scan OCT system

A custom-built spectral-domain point-scan OCT system was employed for both phantom and rodent ORG experiments, as previously reported [35]. The sample arm was modified according to specific imaging requirements: For phantom experiments, a scan lens (LSM04-BB, Thorlabs, USA) was placed after the galvo scanner to focus the beam on the sample, resulting in a spot size (full-width at half-maximum, FWHM) of 19.5 µm in air. For rodent ORG experiments, a scan lens (80 mm doublet) and an ocular lens (30 mm and 25 mm doublet) were used to conjugate the galvo scanner to the pupil plane. Their focal lengths were chosen to achieve smaller beam size and enlarged field of view. The theoretical lateral resolution was estimated to be 7.2 µm (FWHM) based on a standard rat eye model [36].

### 3.2 Phantom fabrication and imaging protocol

Two synthetic phantoms were fabricated by mixing titanium dioxide (TiO2) powder and polydimethylsiloxane (PDMS). Sample #1 had a TiO2-to-PDMS weight ratio of 5% and exhibited fully developed speckles, while a weight ratio of 0.45% was used for Sample #2 to obtain sparsely distributed scatterers. The mixtures were then vacuumed for 20 minutes and cured in an oven (90°C) for 30 minutes. The phantom patch (2 × 2 × 1 mm) was cut off and attached to the surface of a high-precision piezo actuator (P-888.91, Physik Instrumente, Germany). A cover glass, as a static reference, was glued to a lens mount and placed over the phantom without contact. The base of the piezo actuator and the cover glass were connected to the last lens of the OCT sample arm using mechanical frames to reduce the disturbance from environmental vibrations.

To evaluate the performance of subpixel motion correction techniques as introduced in Section 2.4, we designed and implemented the following experiment protocols:

*Protocol 1.* The piezo actuator was driven by a sinusoidal voltage with a frequency of 1 Hz, resulting in a controlled axial vibration of Sample #1 with a peak-to-peak amplitude of ∼2.7 μm. We selected a 1 Hz vibration frequency to minimize distortion within each B-scan. The recording consisted of 600 repeated B-scans, each with 1000 A-lines, acquired at 200 Hz.

*Protocol 2.* The piezo actuator was turned off so that Sample #1 was static, while all other settings were the same as those in Protocol 1.

*Protocol 3.* The same voltage input as in Protocol 1 was used to drive the piezo actuator, resulting in a controlled vibration of Sample #1 in the axial direction with the same frequency. The galvo scanner was turned off. In the recording, a total of 400 repeated A-lines were acquired with a time interval of 5 ms.

*Protocol 4.* To assess motion correction in both axial and lateral directions, we tilted the piezo actuator to create a vibration direction of approximately 70° relative to the OCT beam. Three representative samples were included: two phantoms (Sample #1 and #2), and a cover glass (Sample #3). The piezo actuator was driven by a sinusoidal voltage with a frequency of 1 Hz, resulting in a controlled vibration of ∼8.5 μm in combined directions. The recording consisted of 600 repeated B-scans, each with 1000 A-lines, acquired at 200 Hz.

*Protocol 5.* The piezo actuator was turned off so that the sample remained static. All other settings were identical to those in Protocol 4.

### 3.3 Animal preparation and optoretinogram imaging

The experiments were conducted in compliance with the guidelines and approval from Institutional Animal Care and Use Committee (IACUC), SingHealth (2020/SHS/1574). Eight Brown Norway rats were included in the imaging experiments. The animals were anesthetized with ketamine and xylazine cocktail and fixed in a custom-built stereotaxic holder with integrated bite bar and ear bar to mitigate the eye movements induced by breathing and heartbeat. Before the OCT imaging, the pupil of the animal was dilated with a drop of 1% Tropicamide (Alcon, Geneva, Switzerland) and 2.5% Phenylephrine (Alcon, Geneva, Switzerland). During the imaging, the cornea was kept moist with a balanced salt solution.

In each recording, 800 repeated B-scans were acquired at the same position with each B-scan consisting of 1000 A-lines. Each dataset took a total acquisition time of 4 seconds. To measure the rodent ORG in vivo, visual stimulation was generated by a white light LED (MCWHLP1, Thorlabs, USA). The beam was collimated by an aspheric condenser lens. Then, the stimulation light was converged by the ocular lens (30 mm and 25 mm doublet), resulting in a 43.4 ° Maxwellian illumination on the posterior eye. A short flash with a duration of 2 milliseconds was delivered to the eye 1 second after the recording started.

The phase difference between the inner segment/outer segment junction (IS/OS) and the rod outer segment (ROS) was extracted to evaluate the functional response of photoreceptors to light stimuli. This self-referencing step eliminates undesired phase drift across B-scans. The IS/OS and ROS were segmented using an automatic segmentation algorithm based on graph theory and dynamic programming [37]. In addition, multiply scattered light from superficial large vessels can cause tail artifacts that disturb phase stabilities beneath these vessels [38]. Consequently, the regions below the superficial large vessels were excluded from subsequent phase analysis. To delineate large vessels, a binary OCT angiogram was generated by setting an adaptive threshold on the inverse SNR and complex-valued decorrelation space [39].

### 3.4 Adaptive optics line-scan OCT system for human optoretinogram imaging

A custom-built adaptive optics (AO) line-scan OCT system was constructed for human ORG imaging [11, 40], and the layout can be found in previous report [41]. For human optoretinogram imaging, the experiment was approved by the University of Washington Institutional Review Board and was conducted in compliance with the tenets of the Declaration of Helsinki. Three emmetropic subjects with no known retinal pathologies were enrolled in the study. All subjects signed an informed consent before their participation. Cycloplegia was introduced with Tropicamide 1% ophthalmic solution (Akorn Inc.) before the OCT imaging to dilate the pupil and increase numerical aperture. Prior to recording the human ORG signal, each subject underwent a dark adaption period of 3 to 4 minutes. Three experiment protocols were employed:

*Protocol 1.* 50 volumes (400 B-scans each) were acquired without visual stimulus at a B-scan rate of 6 kHz and a volumetric rate of 12.8 Hz.

*Protocol 2.* A 20-ms visual stimulus (660±10 nm LED in Maxwellian view) was delivered after 10 volumetric scans. 40 volumes were recorded using the same parameters as Protocol 1.

*Protocol 3.* 50 volumes (600 B-scans each) were acquired without visual stimulus at a B-scan rate of 12 kHz and a volumetric rate of 17.0 Hz.

## 4. Results

### 4.1 The numerical validation of motion-induced phase error in FD-OCT, and its accurate correction by PRESIR

To examine the proposed motion-induced phase error, we conducted numerical simulations using a close-to-reality model capable of simulating speckle patterns in FD-OCT [42]. The simulation parameters, including the power spectrum of the light source, lateral resolution, and lateral sampling rate, were set to be consistent with the phantom experiment. As shown in Fig. 1a, we gradually translated the simulated sample in the axial direction by 3 µm with a step size of 0.01 µm. We calculated and unwrapped the phase differences between the subsequent positions and the initial position to obtain the phase change of each pixel. Instead of observing the phase changes in all pixels being consistent with the global translational shifts, we noticed a clear error band that fits the prediction of our theoretical analysis (Fig. 1a). Moreover, as the OCT image gradually decorrelated when we increased the displacement to 3 µm, the phase error band also broadened dramatically. Figure 1b illustrates the distribution of displacements measured from the sample that underwent lateral movement. As expected, the center of the distribution remained at 0, while the increase of lateral shift gradually broadened the distribution.

**Fig. 1.**
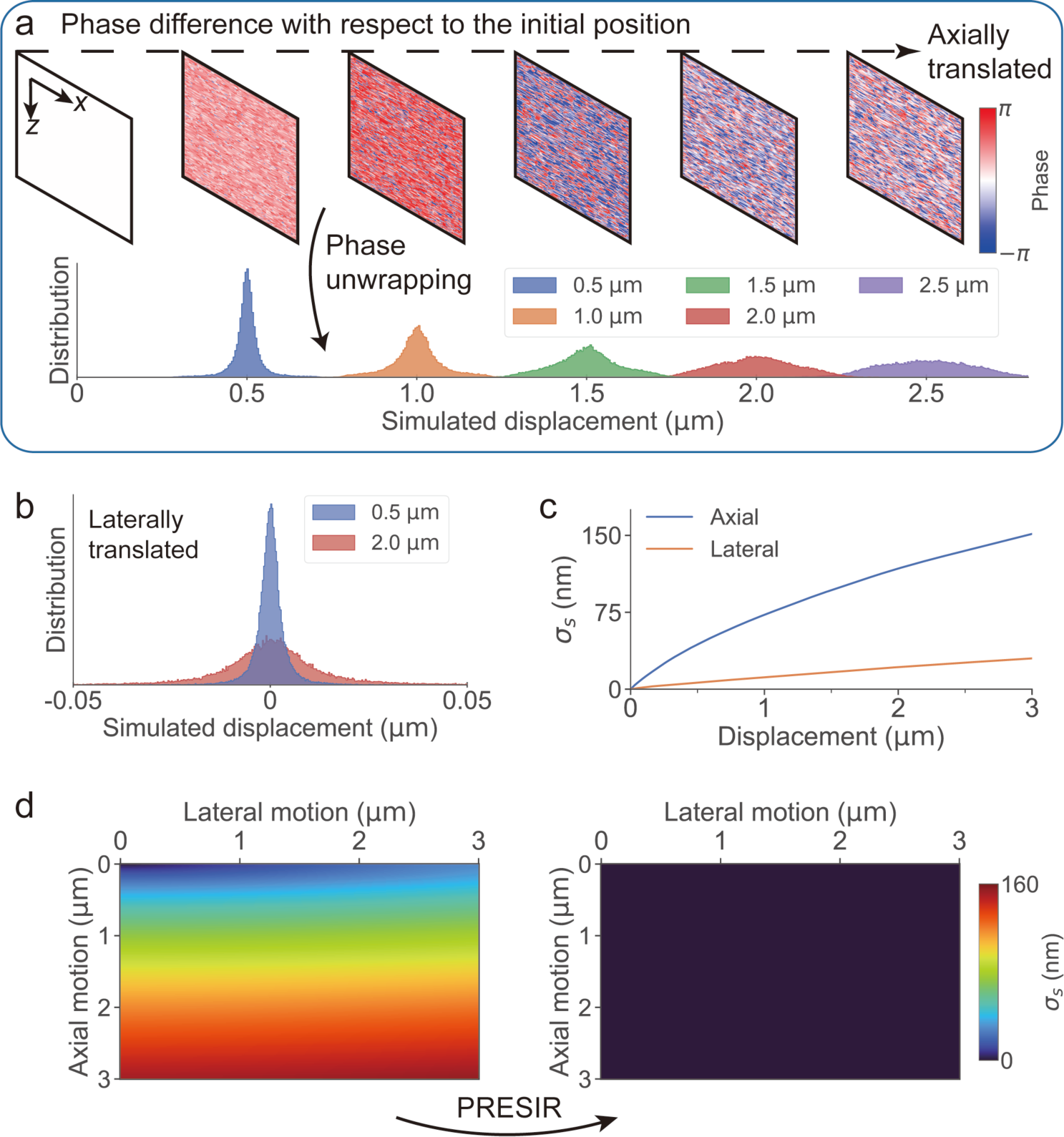
Simulation analysis showing the motion-induced phase error and its accurate correction by PRESIR. (a) Top: The raw phase differences with respect to that at the initial position when moving the simulated sample along the axial direction. Bottom: The distribution of the displacements measured from the above raw phase signals. (b) The distribution of displacements when the sample underwent lateral movement. (c) The motion detection accuracy, calculated from the standard deviation of the phase changes across pixels (*σ*_*s*_), when the sample was translated along the axial or the lateral direction. (d) The motion detection accuracy (left) before and (right) after motion correction using PRESIR when the sample was moved in both axial and lateral directions.

To quantify the motion detection accuracy, we calculated the standard deviation of measured phase changes across pixels for each translation position. Figure 1c shows that the spatial standard deviation of the detected motion (*σ*_*s*_) across all the pixels gradually increased to ∼150 nm and ∼30 nm when the sample was translated by 3 µm along the axial and lateral directions, respectively. This disparity between axial and lateral movements stemmed from the axially compressed OCT PSF. In our simulation and phantom experiments, the lateral resolution (19.5 µm, FWHM) was much lower than the axial resolution (1.9 µm, FWHM), resulting in less significant changes in the PSF-weighted reflectivity (Eq. 5) for lateral motion. Figure 1d also demonstrates that the motion-induced phase error was dominated by axial motion.

Our PRESIR method can restore the original sampling pixels for translationally shifted OCT images, thus restoring the original PSF-weighted reflectivity. As shown in Fig. 1d, *σ*_*s*_measured from the phase signals corrected by the PRESIR method remained near zero.

### 4.2 Error-free motion detection and SNR-limited phase sensitivity validated by moving phantom imaging

To further validate the effectiveness of our PRESIR method in correcting the motion-induced phase error, we conducted phantom imaging experiments. In the first set of experiments (see Section 3.2, Protocols 1-3), we vibrated the sample along the axial direction (Fig. 2), as the simulation analysis indicated that axial movement was the primary contributor to motion-induced phase error considering an axially compressed OCT PSF. In addition, to demonstrate the PRESIR method’s performance in the presence of both axial and lateral motion and its applicability to various samples/scenarios, we conducted a second set of experiments (see Section 3.2, Protocols 4 and 5), where we introduced two-dimensional vibrations on three representative samples (Fig. 3).

**Fig. 2.**
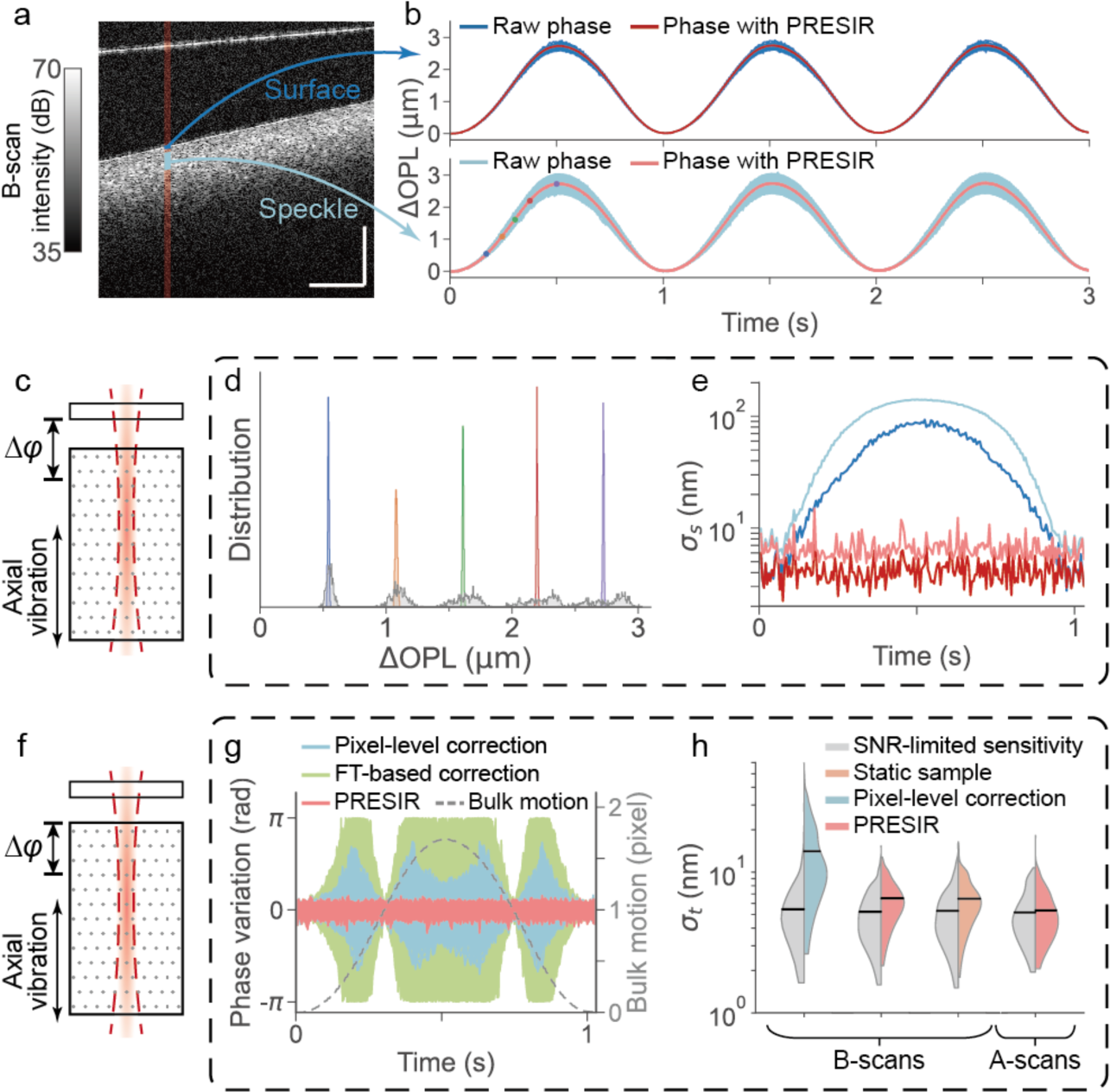
Phantom experiments were conducted to validate the motion-induced phase error and our proposed PRESIR method in the presence of axial bulk motion. (a) The structural OCT image of the phantom’s cross-section. Scale bar: 300 µm. (b) The OPL change measured from the temporal phase change on the phantom surface across 20 adjacent A-lines (blue region in Fig. 2a) and from the internal speckles (cyan region in Fig. 2a). The blue and cyan curves indicate the results without correction, while the red and pink curves denote the results after PRESIR. (c) Schematic of the sample undergoing axial vibration. We extracted the sample’s axial movement by calculating the phase difference between the sample and the cover glass to simulate application scenarios in which the sample’s axial bulk motion is of interest. (d) The distribution of the bulk motion measured from the speckles at 5 time points (as labeled by the colored dots in Fig. 2b). (e) At each time point, the standard deviation of the OPL change across space (*σ*_*s*_) in four configurations (on the surface or from the speckles, with or without PRESIR) is represented by a curve corresponding to the same legend in Fig. 2b. (f) To assess the sensitivity of detecting local deformation during the sample’s axial movement, we calculated the phase differences between the internal speckle patterns and the phantom surface when the sample is under controlled axial vibration. (g) Phase variation at individual pixels in the speckle patterns (the cyan region in Fig. 2a) with respect to the phantom surface after corrected by the pixel-level correction (cyan curves), the FT-based correction (green curves), and the PRESIR (pink curves). The dashed gray line indicates the sample bulk motion. (h) The standard deviation of OPL change over time (*σ*_*t*_) under different conditions (colored halves of the violin plots) and the corresponding theoretical estimated phase sensitivities limited by the SNR (gray halves of the violin plots).

**Fig. 3.**
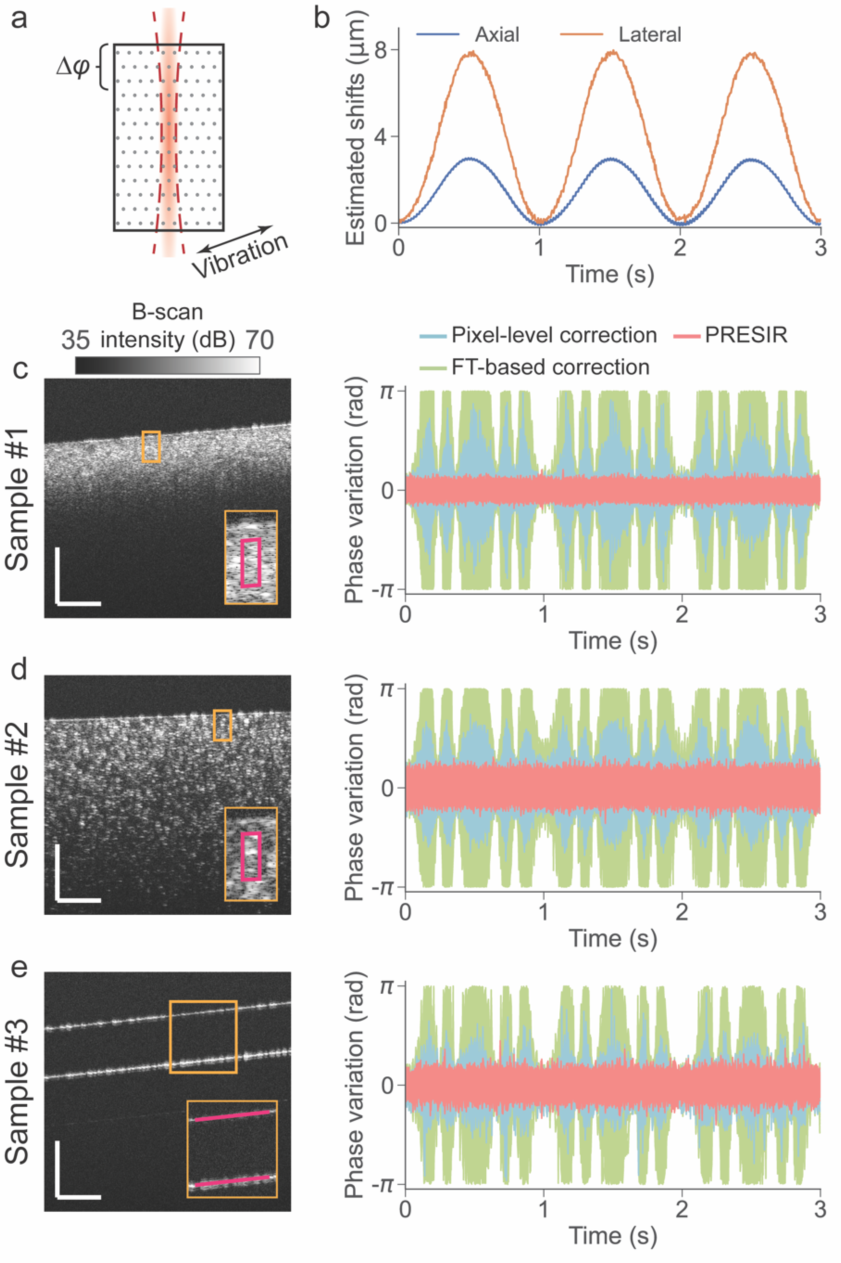
Phantom experiments demonstrating the effectiveness of the proposed PRESIR method in correcting combined axial and lateral bulk motion. (a) Schematic of a sample undergoing two-dimensional vibration. Phase stability was evaluated by calculating the phase difference between speckle patterns (regions enclosed by magenta outlines in Figs. 3c and 3d) and the phantom surface, or between the two surfaces of the cover glass (magenta lines in Fig. 3e). (b) A representative bulk motion measured from Sample #1 in (c). (c)-(e) Left: Structural images of three representative samples (Samples #1, #2, and #3) described in Section 3.2. Scale bar: 300 μm. Right: Phase variations measured at individual pixels after applying pixel-level correction (cyan curves), FT-based correction (green curves), and PRESIR (pink curves).

As depicted in Fig. 2a, a synthetic phantom (refer to Section 3.2) was attached to a piezo actuator driven by a sinusoidal voltage input to mimic a moving biological tissue; a cover glass served as a static reference to eliminate undesired phase drift. Temporal phase changes of individual pixels, measured from the phantom surface (blue curves in Fig. 2b) and the internal speckles (cyan curves in Fig. 2b, with the outliers removed as per the criterion in Appendix C) exhibited similar sinusoidal patterns due to axial motion. Meanwhile, noticeable motion-dependent deviations were observed from the phase changes of individual pixels (blue and cyan curves in Fig. 2b), likely due to the motion-induced phase error discussed in the preceding theoretical analysis. Supplementary Video 1 reveals the spatiotemporal evolution of phase errors measured from internal speckle patterns (an area around the cyan region in Fig. 2a). The phase errors in individual pixels were repetitive over periodic sinusoidal bulk tissue motion, which corroborated our theoretical analysis of motion-induced phase errors as deterministic errors instead of stochastic noise.

To correct the motion-induced phase error, we estimated the sample displacement by calculating the average phase change from the phantom surface. After applying the PRESIR method for motion correction and removing the outliers (refer to Appendix C for the criterion), we observed a significant reduction in motion-dependent errors, resulting in consistent OPL changes across all surface pixels (red curves in Fig. 2b) and internal speckle patterns (pink curves in Fig. 2b). Fig. 2d shows the distribution of the motion measured from the speckles at five time points (as shown by the colored dots in Fig. 2b). Compared with the raw phase results (the bottom gray areas in Fig. 2d), the PRESIR method achieved much sharper measurement distributions (the colored peaks in Fig. 2d). After motion correction using PRESIR, the standard deviation of the detected OPL change across individual pixels (*σ*_*s*_) was reduced to a level no longer dependent on motion (red and pink curves in Fig. 2e).

An important application of the proposed PRESIR method is for improving the phase sensitivity during the measurement of the nanoscopic local deformations in moving biological tissues. To evaluate phase sensitivity/stability, we measured the phase differences of individual pixels in the speckle patterns with respect to the phantom surface (Fig. 2f). Figure 2g illustrates phase variations over time in the cyan regions of Fig. 2a, with outliers removed according to the criterion in Appendix C. When using conventional pixel-level correction, the phase error (cyan curves in Fig. 2g) increased gradually as the residual bulk motion, defined as the deviation of the bulk motion (dashed gray curve in Fig. 2g) from the nearest integer, approached half a pixel. Meanwhile, the FT-based method led to even larger phase error (green curves in Fig. 2g). In contrast, after applying the PRESIR method for motion correction, phase variation across all pixels was significantly reduced (pink curves in Fig. 2g), with the remaining phase variation becoming independent of the sample’s bulk motion.

In order to quantify the phase sensitivities at individual pixels, we analyzed the temporal standard deviation (*σ*_*t*_) of the OPL variation over 3-second recordings. Additionally, we calculated the theoretical SNR-limited phase sensitivity for each pixel [2]. As shown in Fig. 2h, due to the sample movement, the phase variations after pixel-level correction (16.4 ± 8.9 nm, mean ± standard deviation) were significantly higher than the theoretical SNR-limited phase sensitivities (6.2±3.8 nm). Motion correction using the PRESIR method improved the phase stabilities of individual pixels across repeated B-scans to 6.7±2.2 nm. Meanwhile, the SNR-limited phase sensitivities were adjusted to 5.5±2.2 nm owing to the stabilized SNRs. Such performance closely resembled results measured from a static sample across repeated B-scans (6.7±2.3 nm, with SNR-limited phase sensitivities at 5.6±2.4 nm). We determined that the remaining deviation from the SNR-limited phase sensitivity was caused by the instability of the galvo scanner during the scanning [3]. Once we turned off the galvo scanner for repeated A-scans, the phase stabilities measured after motion correction by PRESIR from eight independent trials (5.7±2.2 nm) were considerably close to the SNR-limited phase stability (5.6±2.1 nm). These results demonstrate that the PRESIR method effectively eliminates motion-induced phase errors in phase-sensitive OCT, achieving the fundamental phase sensitivity limit determined by the SNR.

In the presence of combined axial and lateral bulk motion (Fig. 3a), we adopted the single-step DFT approach (see Methods) to estimate the bulk tissue motion (Fig. 3b). To evaluate the phase stability in Samples #1 and #2 (PDMS samples as described in Section 3.2), we calculated the phase difference between the internal speckle patterns (regions enclosed by magenta outlines in Figs. 3c and 3d) and the phantom surface. Outliers were removed according to the criterion described in Appendix C. For Sample #3 (cover glass), we calculated the phase difference between two surfaces to assess the phase stability (magenta lines in Fig. 3e). The pixel-level and FT-based methods resulted in phase variations dependent on the bulk motion. In comparison, our proposed PRESIR method eliminated the motion-dependent phase variations. A quantitative comparison using the temporal standard deviation (*σ*_*t*_) of the OPL variations (Table 1) demonstrated that the phase stability achieved with PRESIR surpassed that of the benchmark pixel-level correction and was comparable to a static sample.

**Table 1:**
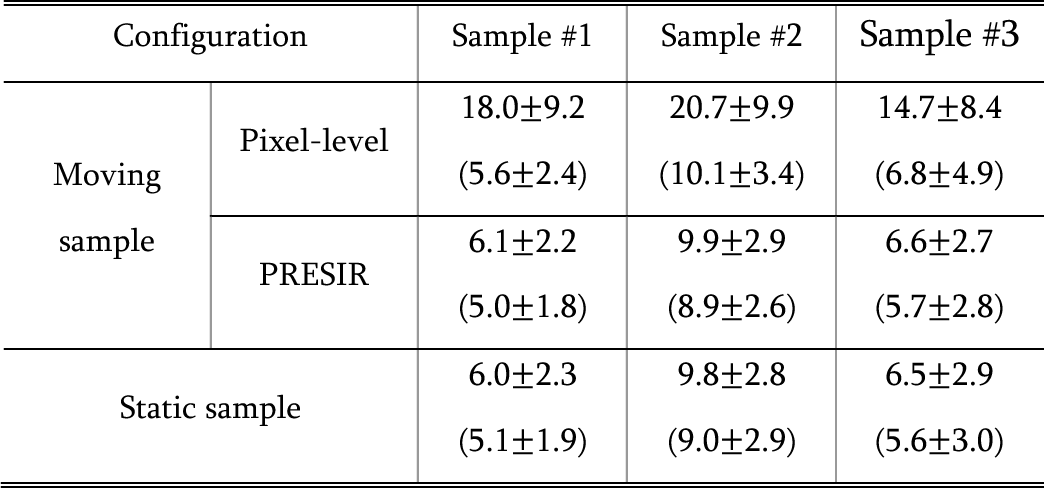
Measured phase stabilities (*σ*_*t*_) and estimated SNR-dependent phase stabilities (in brackets) from three representative samples using different configurations. All values are expressed in nm as mean±standard deviation.

| Configuration |  | Sample #1 | Sample #2 | Sample #3 |
| --- | --- | --- | --- | --- |
| Moving sample | Pixel-level | 18.0±9.2<br>(5.6±2.4) | 20.7±9.9<br>(10.1±3.4) | 14.7±8.4<br>(6.8±4.9) |
| | PRESIR | 6.1±2.2<br>(5.0±1.8) | 9.9±2.9<br>(8.9±2.6) | 6.6±2.7<br>(5.7±2.8) |
| Static sample | | 6.0±2.3<br>(5.1±1.9) | 9.8±2.8<br>(9.0±2.9) | 6.5±2.9<br>(5.6±3.0) |

### 4.3 Improved motion detection sensitivity for in-vivo label-free tissue dynamics imaging at the nanoscopic scale

The phantom imaging experiment demonstrated the PRESIR’s ability to eliminate the motion-induced phase error when measuring the sample’s bulk motion (Figs. 2c-e). The experiment also revealed the significantly improved phase sensitivity/stability when the sample’s local deformation is of interest (Figs. 2f-h and Fig. 3). To further investigate motion detection performance during label-free imaging of nanoscopic tissue dynamics in vivo, we conducted ORG imaging experiments in both rodents and humans.

For rodent ORG imaging, we performed repeated B-scans in a wild-type rat’s retina using a point-scan OCT system. Fig. 4a shows a structural image and a time-elapsed M-scan at one A-line. As shown in Fig. 4b, we estimated the bulk motion of the retina using the efficient single-step DFT approach (see Methods). The PRESIR method corrected the apparent bulk motion in the raw M- scan (Fig. 4c, 1st column), resulting in nearly static retinal layers (Fig. 4c, 4th column). In contrast, pixel-level correction left serrated residual motion (Fig. 4c, 2nd column), while the FT-based correction method introduced severe side lobes unexpectedly (Fig. 4c, 3rd column).

**Fig. 4.**
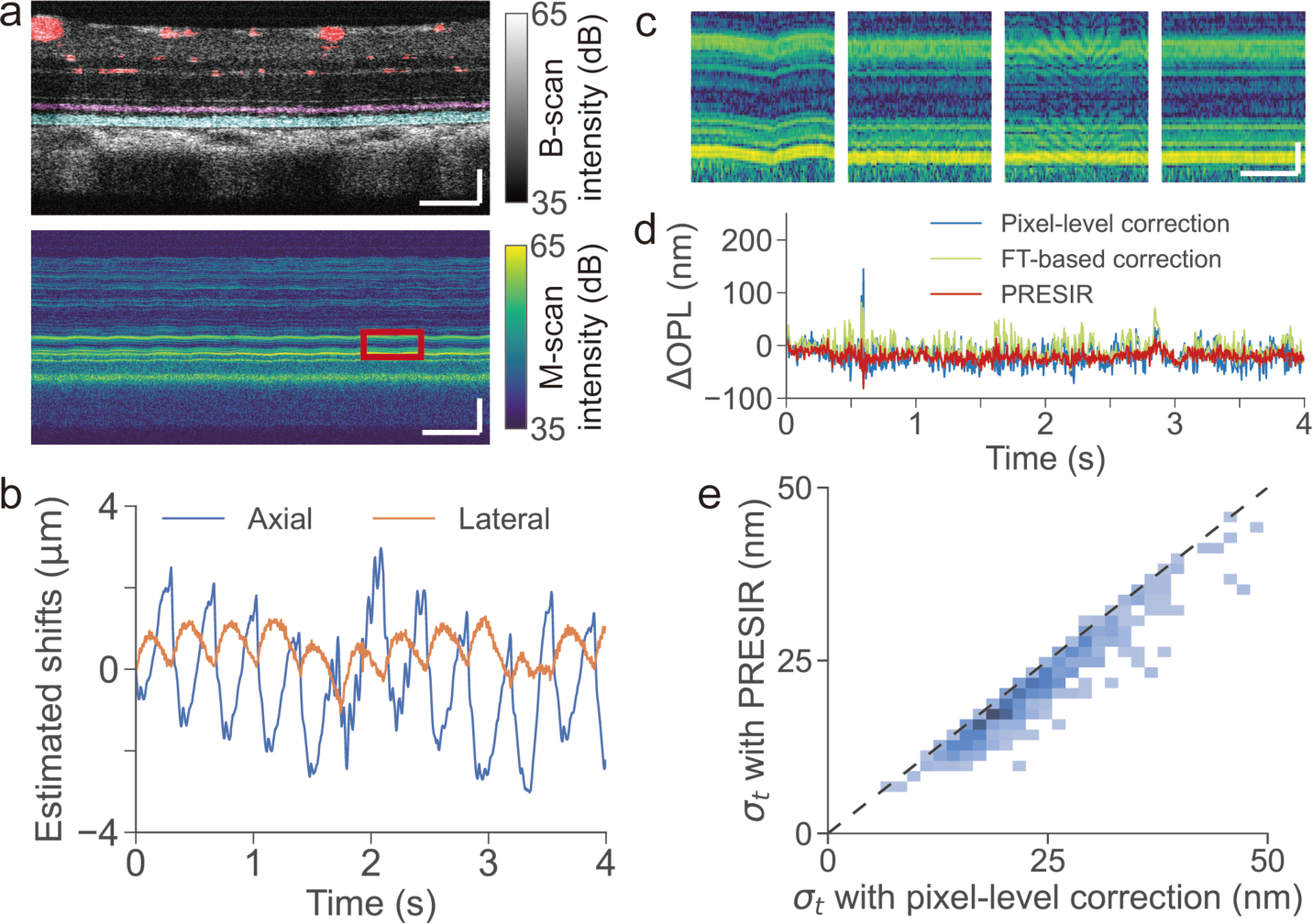
In-vivo rodent retinal imaging using a non-AO point-scan OCT. (a) Top: Structural image of a Brown Norway rat’s retina. The pink, cyan, and red areas represent the inner segment/outer segment junction (IS/OS), the rod outer segment (ROS) and blood vessels, respectively (refer to Section 3.3). Bottom: a representative time-elapsed M-scan at one A-line. Scale bar in spatial dimension: 100 µm. Scale bar in temporal dimension: 0.5 s. (b) Bulk motion of the retinal tissue estimated by the efficient single-step DFT approach. (c) Enlarged M-scans (enclosed red window in Fig. 3a) without correction (1st column), after pixel-level correction (2nd column), FT-based method correction (3rd column), and PRESIR method correction (4th column). Scale bar in spatial dimension: 20 µm. Scale bar in temporal dimension: 0.2 s. (d) The phase differences between the brightest pixel in IS/OS and ROS on one representative A- line after the pixel-level correction (blue line), the FT-based correction (green line), and the PRESIR (red line), when no light stimulus was delivered to the retina. (e) Standard deviation distribution of phase fluctuation between IS/OS and ROS over time (*σ*_*t*_) across A-lines, comparing pixel-level correction to PRESIR. The dashed black line represents the barrier where the two methods yield the same performance.

Using the IS/OS layer as a reference to cancel out phase drifts between B-scans, Supplementary Videos 2 and 3 demonstrate the pulsatile deformation before and after applying the PRESIR method. Compared to raw phase signals, the PRESIR method effectively eliminated motion-induced phase errors and enabled reliable visualization of pulsatile deformation, highlighting the importance of optimal bulk tissue correction in detecting local tissue deformation. To quantitatively compare the phase stability enabled by different motion correction methods, the phase difference between the brightest pixels in the IS/OS (the pink area in Fig. 4a) and the ROS (the cyan area in Fig. 4a) was extracted from individual A-lines (Fig. 4d). Across the entire field of view, except for the region below large superficial vessels, the standard deviation of such phase differences over time (*σ*_*t*_) was reduced to 21.1±7.1 nm (mean±standard deviation) by PRESIR, compared with 23.8 ± 7.4 nm after pixel-level correction and 23.2 ± 8.0 nm after FT-based correction. Fig. 4e demonstrates that the PRESIR method significantly improved the phase stability compared with pixel-level correction.

Figure 5a, in its first column, presents a violin plot comparing the phase stabilities previously shown in Fig. 4e. To assess the robustness of our proposed method, we extended the validation to an additional seven rats. Across all eight animals (Fig. 5a), our proposed method achieved a phase stability of 21.3±8.0 nm, outperforming both the pixel-level correction (24.3±8.5 nm) and the FT-based correction (23.5±8.8 nm). In ORG measurements with a 2-ms light stimulus delivered at the time *t* = 0, the PRESIR method produced more stable and less noisy ORG signals than pixel-level correction (Fig. 5b), in which case a clean ORG signal after averaging across 6 pixels is shown in Fig. 5c.

**Fig. 5.**
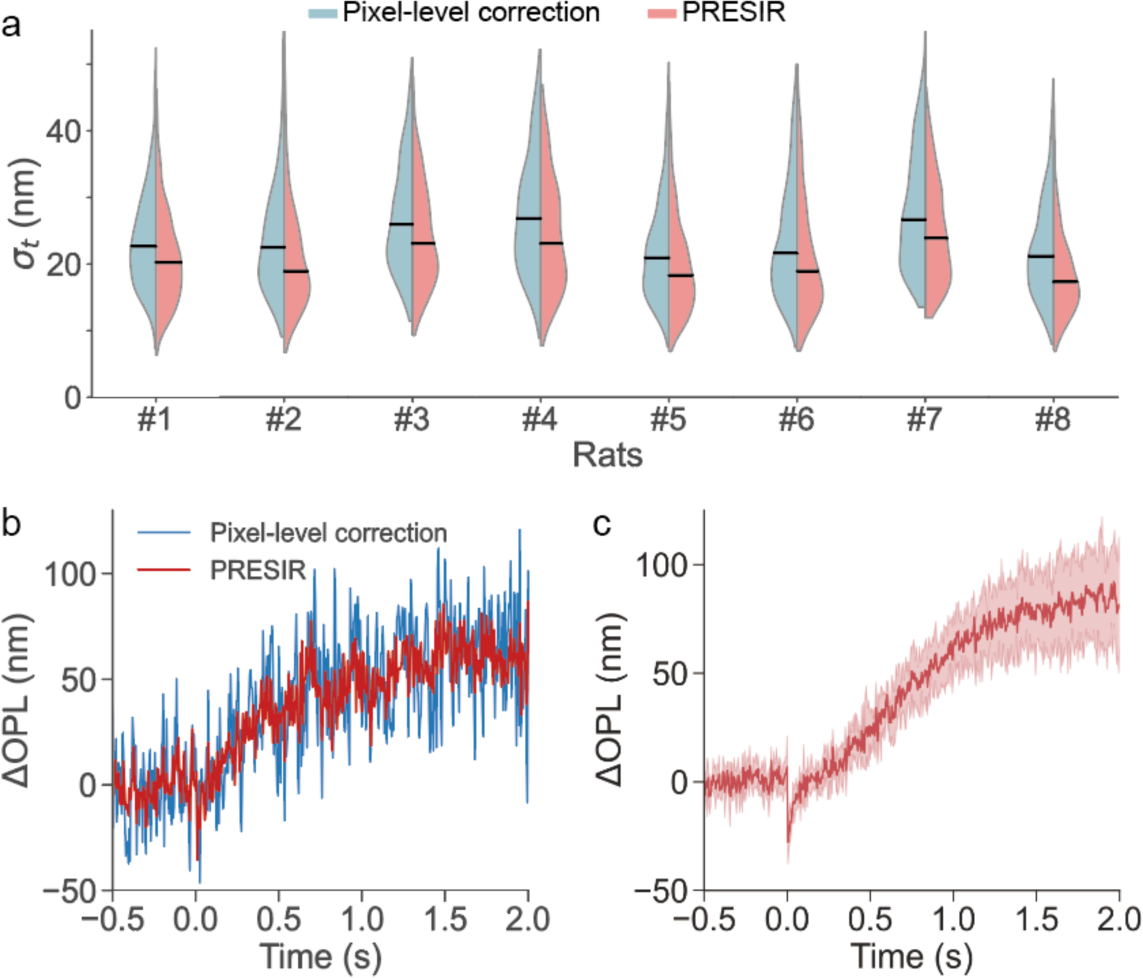
Phase stabilities measured from eight rodents and representative ORG signals in response to visual stimuli. (a) Temporal phase stabilities measured from eight rodents after the pixel-level correction and PRESIR method. (b) Representative ORG signals after the pixel-level correction (blue line) and the PRESIR (red line) obtained from one A-line without averaging. (c) ORG signal of a wild-type rat after the PRESIR. The red line represents the average of 6 phase signals with high phase stability during the pre-stimulus period (pre-stimulus *σ*_*t*_ < 10 nm), and the color band denotes the range of their standard deviation.

We further tested the proposed method for enhancing motion detection performance in human ORG imaging using a high-speed adaptive-optics line-scan OCT system. Figs. 6a and 6b show the en-face image at the cone outer segment tips (COST) and a structural image at one cross-section of the retina. The translational bulk motion of each volume was estimated using a modified coarse-to-fine approach (refer to Methods). Regarding motion correction performance, the PRESIR method achieved more stable retinal layers (Fig. 6c, right) compared with pixel-level correction (Fig. 6c, left) and the FT-based method (Fig. 6c, middle). Fig. 6d shows the representative phase differences between the IS/OS and the COST at two individual A-lines without light stimulus following Section 3.4, Protocol 1. The phase stability was evaluated by calculating the standard deviation of the phase difference over time (*σ*_*t*_). As shown in Fig. 6e, across the entire field of view while excluding gaps between cones based on averaged intensity from IS/OS and COST layers, the PRESIR method achieved higher phase stability with smaller phase fluctuation standard deviations (15.3 ± 9.0 nm, mean ± standard deviation) compared with pixel-level correction (19.7±10.0 nm). In contrast, the FT-based correction significantly degraded the phase stability (46.4±17.3 nm). Interestingly, our PRESIR approach not only suppressed unwanted fluctuations (Fig. 6f, location 2) but also recovered ORG signals (obtained via Section 3.4, Protocol 2) that would otherwise suffer from phase unwrapping errors when using the pixel-level correction method (Fig. 6f, location 1).

**Fig. 6.**
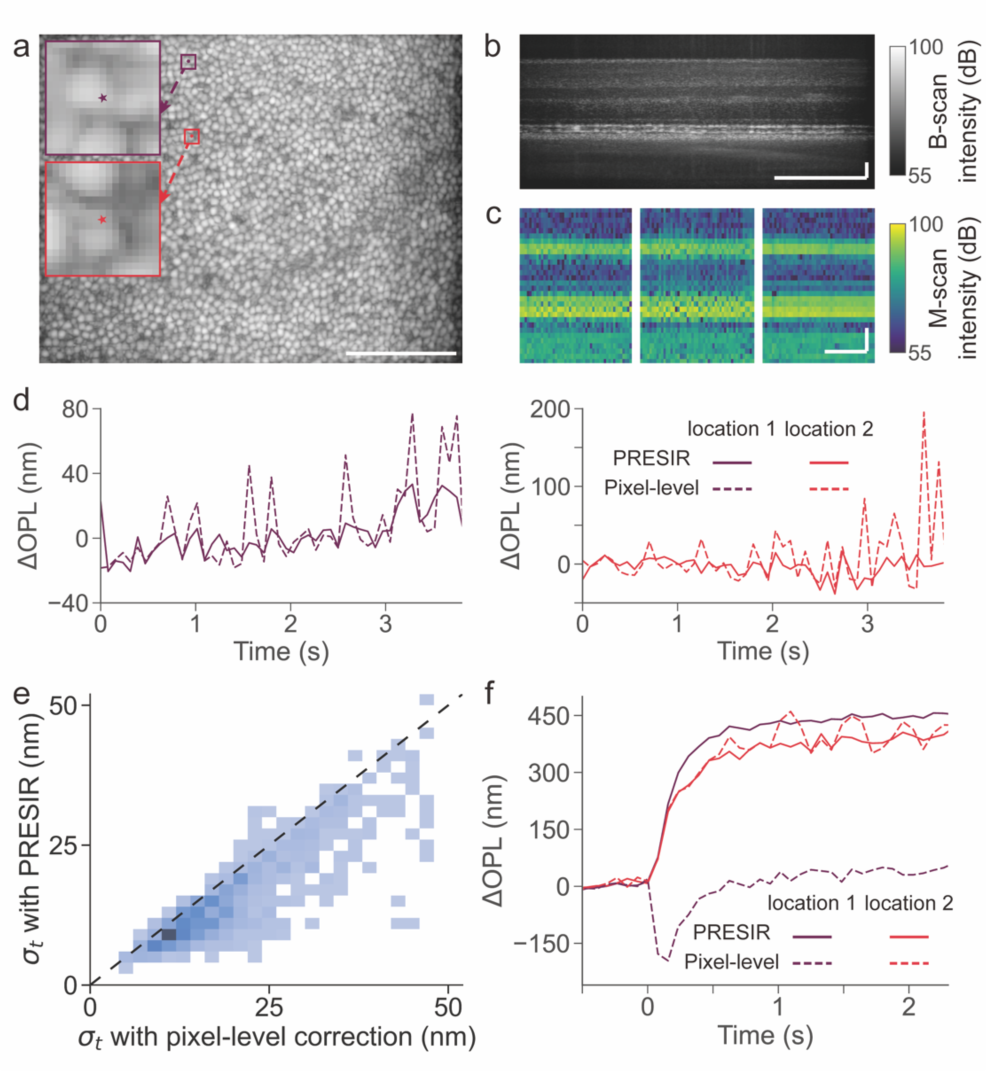
In-vivo human optoretinography using an AO line-scan OCT. (a) En-face image of the cone outer segment tips (COST) layer in a human retina. (b) Cross-sectional structural image of the same retina. Scale bar: 100 µm. (c) Representative time-elapsed M-scans corrected by the pixel-level method (left), the FT-based method (middle) and the PRESIR method (right). Scale bar in spatial dimension: 20 µm. Scale bar in temporal dimension: 1 s. (d) Representative phase differences between the IS/OS and COST at individual A-lines after the pixel-level correction (dashed lines), and the PRESIR (solid lines), when no light stimulus was delivered to the retina. (e) The standard deviation distribution illustrates phase fluctuations between the IS/OS and COST over time (*σ*_*t*_) across A-lines, comparing pixel-level correction with PRESIR. The dashed black line indicates the point at which the two methods exhibit equivalent performance. (f) Representative human ORG signals measured from individual A-lines (labeled by the dots in Fig. 6a) showing higher phase stability after the PRESIR (solid lines) than those corrected by the pixel-level correction (dash lines).

To assess the generalizability of our findings, we compared the performance of PRESIR with conventional approaches in two additional human subjects using a faster imaging protocol (Section 3.4, Protocol 3). PRESIR consistently demonstrated superior phase stability across both subjects. In Subject #2, PRESIR achieved a phase stability of 12.4±4.3 nm, compared to 18.3±7.2 nm for the pixel-level correction and 50.6±22.2 nm for the FT-based correction (Figs. 7a and 7b). Similarly, in Subject #3, the phase stability after PRESIR was 14.1±5.8 nm, compared to 20.2±8.7 nm after the pixel-level correction and 46.8±18.8 nm after the FT-based correction (Figs. 7c and 7d).

**Fig. 7.**
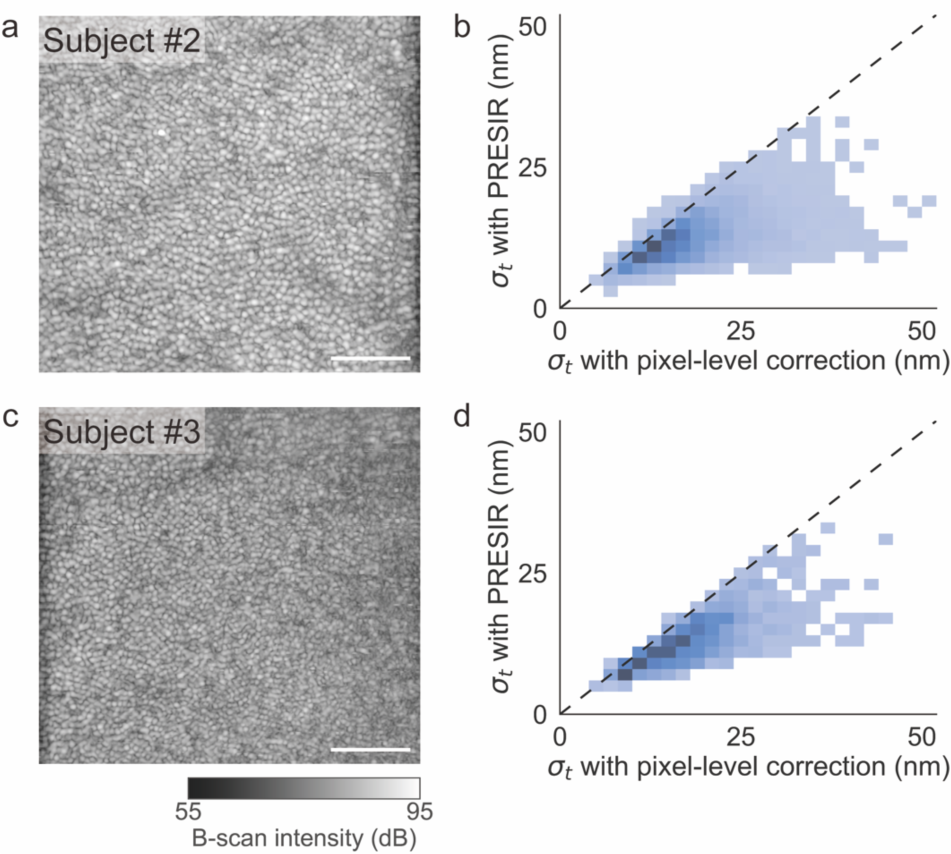
In-vivo human retinal imaging in two additional human subjects. (a) and (c) En-face images of the cone outer segment tips (COST) layer. (b) and (d) Corresponding phase stabilities obtained after the pixel-level correction and PRESIR.

## 5. Discussion

In this article, we present a phase-restoring subpixel image registration (PRESIR) approach for post-hoc image registration in FD-OCT, which allows for translational shifting of OCT images by arbitrary displacements while accurately restoring physically meaningful phase components. We discovered that in moving samples, the phase difference measured by FD-OCT includes both the anticipated OPL change corresponding to the sample movement and a motion-induced phase error arising from alterations in PSF-weighted reflectivity of scatterers. Correcting such phase error requires reproducing the same sampling points at each frame in the time-elapsed recording. By employing the PRESIR method, we achieved phase-sensitive imaging of a moving phantom with a sensitivity close to the fundamental limit set by the SNR. Moreover, we showed that the PRESIR method substantially improved motion detection performance in label-free imaging of nanoscopic tissue dynamics in vivo, particularly in the context of emerging functional assessments using optoretinography.

The residual motion-induced phase error after PRESIR primarily depends on the accuracy of translational motion estimation, which could be further improved by adopting more sophisticated subpixel-level motion estimation methods based on model fitting or optimization strategies [24]. Recently, convolutional neural networks (CNNs) have been used to estimate 3D translational motion between repeated volumetric scans, enabling improved accuracy and reduced computational time [43, 44].

The present study has several limitations, particularly concerning repeated B-scans in rodent ORG experiments, where the out-of-plane motion remained uncorrected. Depending on its magnitude, the out-of-plane motion may decorrelate the speckle patterns and degrade the phase sensitivity. While translational displacements are the primary source of motion artifacts in high-speed OCT imaging [26], motion artifacts due to other types of movements may also exist. Recent studies have employed affine transformation to rectify translation, rotation, and scaling errors [45, 46], while non-rigid B-spline transformation has been used to assess local deformation between repeated volumetric scans [47]. Our proposed method may inspire new subpixel-level phase-restoring motion correction approaches for these scenarios. Meanwhile, our current model assumes a collimated OCT beam, justified by the low numerical aperture typically used in ocular imaging OCT systems to maintain a relatively consistent transverse resolution throughout the entire axial (depth) scan [48]. Additional impacts caused by defocus can be found in Appendix D.

Our proposed PRESIR approach has many potential applications in various OCT imaging modalities. In Doppler OCT, the phase difference between repeated or adjacent densely sampled A-scans is calculated to quantify flow velocity [6]. However, as shown in this study, uncorrected motion-induced phase errors can compromise measurement accuracy even when the tissue movement is as small as 10 nm in the axial direction. In optical coherence elastography, both the speckle-tracking and phase-sensitive detection methods were used to estimate sample displacements [7]. The PRESIR method’s ability to manipulate OCT images as if the sample were physically shifted might enable new motion estimation methods through optimization. In OCT angiography, complex-value-based algorithms offer higher motion contrast, but their performance is susceptible to phase error due to bulk motion [49], which could be effectively suppressed by the PRESIR method. Furthermore, our proposed approach may also benefit computational imaging technologies [50], where phase stability is critical to their in-vivo implementations.

## Acknowledgments

We thank Dr. G. Barbastathis at MIT, Dr. D. Palanker at Stanford University, Drs. A. Roorda and F. Feroldi at the University of California, Berkeley, and Dr. E. Pugh at the University of California, Davis, for fruitful discussions.

## Author contributions

H.L. and T.L. conceived and designed the study. B.T. and L.S. built the ultrahigh-resolution point-scan OCT setup. V.P.P. and R.S. built the adaptive optics line-scan OCT setup. H.L., B.T., and V.P.P. conducted the experiments. V.A.B. supported the animal preparation. H.L. and T.L. analyzed the data and wrote the article. All authors reviewed and edited the article. T.L. supervised the study.

## Data availability

All data needed to evaluate the conclusions in the paper are present in the paper. Additional data related to this paper may be requested from the authors.

## Appendix

### A. FD-OCT signal manifestations in the k-domain, spatial domain, and spatial frequency domain

We assume that individual scatterers along an A-line in the sample follow a reflectivity distribution *η*(*z*), which can be written as,

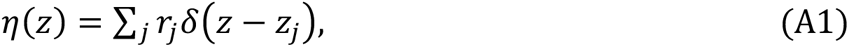

where *z* represents the optical path length (OPL) - defined as the integral of refractive index along the physical depth, *r*_*j*_ stands for the electric field reflectivity of the *j*^th^ scatterer, *z*_*j*_ indicates the axial location of the *j*^th^ scatterer in OPL, and *δ*() denotes the Dirac delta function.

#### 1) Spectral interferogram formed on the detector (k domain)

Under the FD-OCT framework, the corresponding spectral interferogram *Ĩ*(*k*) can be modeled as the linear superposition of scattered signals from each scatterer [18, 42],

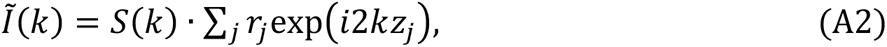

where *k* is the wavenumber and *S*(*k*) is the power spectrum of the light source. For simplicity, we have neglected DC terms, auto-correlation terms, complex conjugate artifacts, and constants like the responsivity of the detector.

If we further take the lateral direction (*x*) into consideration and assume an illumination beam with a Gaussian profile, the field reflectivity map *η*(*x*, *z*) and its corresponding two-dimensional (2D) spectral interferogram *Ĩ*(*x*, *k*) can be modeled as [19],

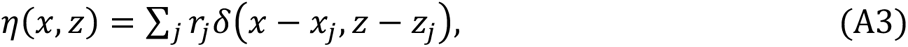

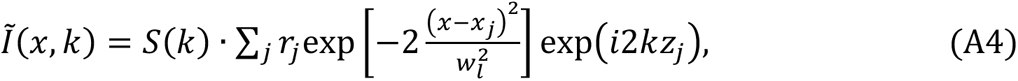

where *x*_*j*_ is the lateral position of the *j*^th^ scatterer, *w*_*l*_ is the 1⁄e^2^ spot radius of the OCT beam focused on the sample.

#### 2) Complex-valued OCT image obtained from a Gaussian-shaped illumination spectrum (spatial domain)

The depth-resolved OCT signal can be reconstructed by performing a Fourier transform along the spectrum (*k*) direction [18],

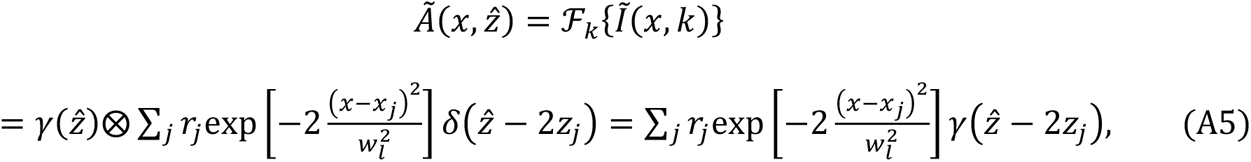

where *Ã* is the complex-valued OCT image, and *ẑ* = 2*z* denotes the round-trip OPL variable. *γ*(*ẑ*) is the coherence function of the OCT system, defined as the Fourier transform of the power spectrum, while ⊗ denotes the convolution operation.

Given a normalized Gaussian-shaped spectrum *S*(*k*), the coherence function can be obtained by,

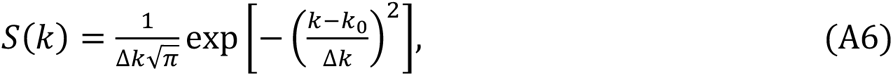

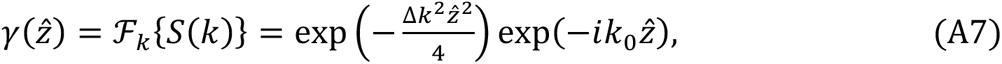

where *k*_0_ is the center wavenumber of the light source spectrum and Δ*k* is the half-width of the spectrum at 1⁄e of its maximum.

Combining Eqs. (A5) and (A7), and substituting *ẑ* by 2*z*, the OCT image can be written as,

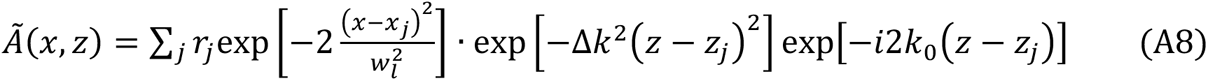

and it can be further simplified as,

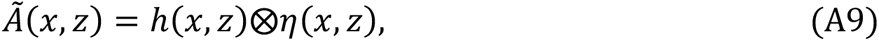

where 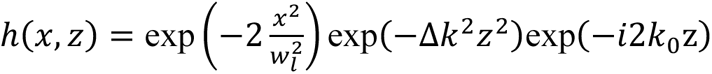 is the point spread function (PSF) of the OCT system.

#### 3) Fourier transform of the complex-valued OCT image (spatial frequency domain)

The Fourier transform of the complex-valued OCT image is,

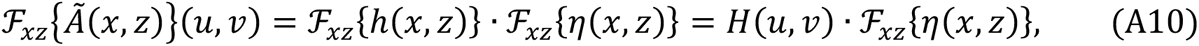

where *u* and *v* represent the lateral and axial spatial frequency. Equation (A10) indicates that the profile of the spatial frequency signal of a complex-valued OCT image depends on the system’s optical transfer function *H*(*u*, *v*), as shown in Eq. (A11),

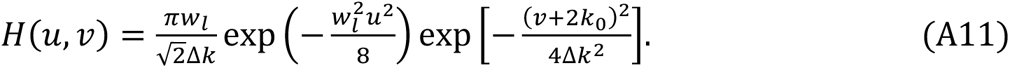

### B. Change in PSF-weighted reflectivity during sample movement

Following Eqs. (1) and (2) in the main text and Eq. (A8) in Appendix A, the complex-valued OCT images 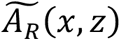 and 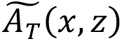 measured from the reference frame and the target frame can be modeled as

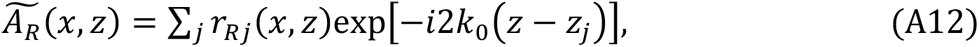

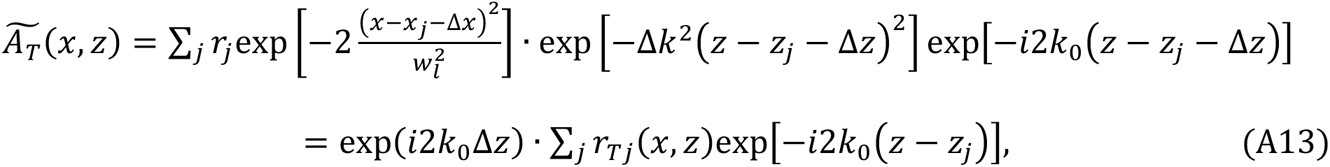

where *r*_*R*j_(*x*, *z*) and *r*_*T*j_(*x*, *z*) are defined as the PSF-weighted reflectivities of scatterers in the reference and target frames:

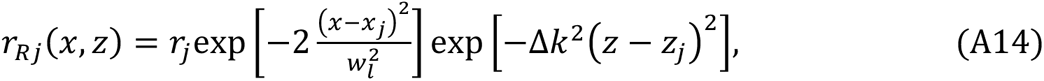

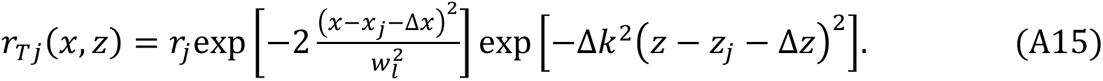

### C. Criterion to remove outliers in phantom experiment

Phase unwrapping errors occurring at a few pixels can distort statistical estimations, causing them to deviate from the actual distributions of the measured phase signals. For example, when calculating the standard deviation of the measured phase signals across a series of pixels, these outliers can result in a substantially overestimated standard deviation, which hinders a reliable estimation of the measurement accuracy. To exclude the outliers, we adopted the criterion introduced in previous studies, which sets the upper and lower thresholds as *q*_3_ + *w*(*q*_3_ – *q*_1_) and *q*_1_ − *w*(*q*_3_ – *q*_1_), where *w* was set to 2 in this study, *q*_1_ and *q*_3_ represent the 25th and 75th percentiles of the sample data, respectively [51]. Assuming a Gaussian distribution of motion-induced phase errors across speckle patterns, we expect this threshold to exclude only 0.2% of the signals. Consequently, the chosen threshold should have a minor impact on the overall signal distribution while effectively suppressing outliers that deviate significantly from the distribution profile.

### D. Impact of defocus on motion correction

Considering a focused Gaussian beam, the spectral interferogram of the reference frame 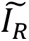 can be written as [48, 52],

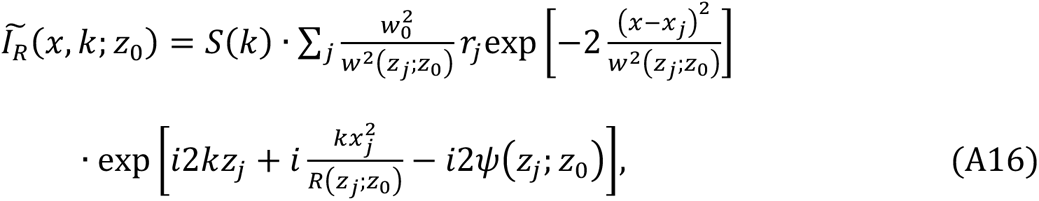

where *z*_0_is the depth position of the focus and *w*_0_is the beam radius at the focal plane. *w*(*z*; *z*_0_) and *R*(*z*; *z*_0_) account for the depth-dependent beam waist size and the phase curvature of the wavefront, respectively. *ψ*(*z*_*j*_; *z*_0_) is the Guoy phase shift. These parameters are given by,

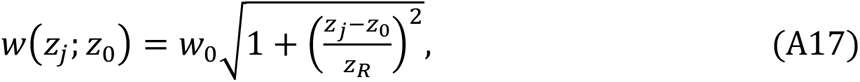

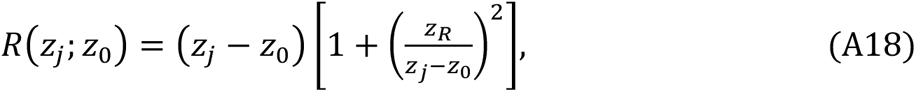

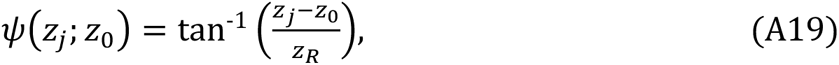

where *z*_*R*_ is the Rayleigh range.

The consistency of the PSF across the lateral direction at a given depth suggests that it should not impact the effectiveness of our lateral motion correction. For simplicity, we will focus on axial displacement in the following derivation to examine the influence of depth-dependent PSF on the performance of axial motion correction. When the sample undergoes a bulk motion of Δ*z* along the axial direction, the corresponding interferogram of the target frame 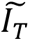 can be written as,

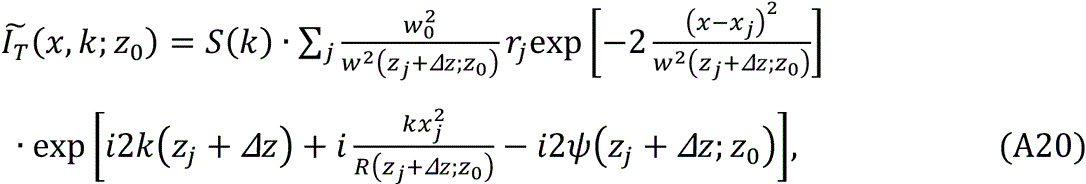

If we conduct the same axial motion correction by multiplying an exponential term exp(−*i*2*k*Δ*z*) as in Eq. (10), the corrected spectral interferogram 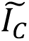 is given by,

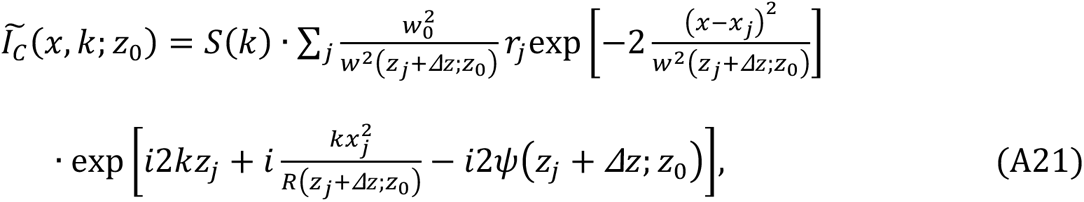

According to Eqs. (A17)-(A19), we have *w*(*z*_*j*_ + Δ*z*; *z*_0_) = *w*(*z*_*j*_; *z*_0_ − Δ*z*), *R*(*z*_*j*_ + Δ*z*; *z*_0_) = *R*(*z*_*j*_; *z*_0_ − Δ*z*) and *ψ*(*z*_*j*_ + Δ*z*; *z*_0_) = *ψ*(*z*_*j*_; *z*_0_ − Δ*z*). By substituting these equations into Eq. (A21), it can be rewritten as,

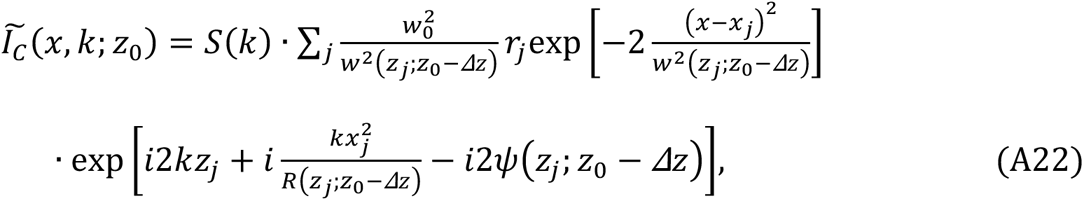

Comparing Eq. (A16) and Eq. (A22), we can conclude that,

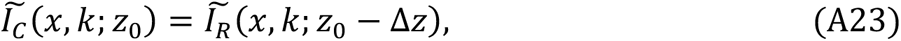

This equation suggests that the corrected frame is equivalent to fixing the sample and shifting the focal plane by −Δ*z*. Given that OCT systems typically employ lenses with a low numerical aperture [48], the impact of focal plane shifts should be minimal, provided that the sample’s bulk movement Δ*z* is significantly smaller than the Rayleigh range (∼200 μm for our rodent retinal imaging). Nevertheless, when the magnitude of bulk tissue motion approaches the Rayleigh range, the accuracy of the motion correction can be adversely affected. In such cases, numerical focusing methods could be implemented to mitigate additional impacts caused by the depth-dependent PSF [53]. Future studies may explore experimental validation of this theoretical analysis.

